# The Spectral Fingerprint of Sleep Problems in Post-Traumatic Stress Disorder

**DOI:** 10.1101/209452

**Authors:** M. de Boer, M.J. Nijdam, R.A. Jongedijk, Olff, W.F. M. Hofman, L.M. Talamini

**Affiliations:** Brain and Cognition, Dept. of Psychology, University of Amsterdam, Nieuwe Achtergracht 129 B, 1018 WS Amsterdam; UvA – Amsterdam Brain and Cognition; Center for Psychological Trauma, Department of Psychiatry, Academic Medical Center at the University of Amsterdam, Amsterdam, The Netherlands; Foundation Centrum’45, partner in Arq, Oegstgeest, The Netherlands; Arq Psychotrauma Expert Group, Diemen, The Netherlands

**Keywords:** Post-traumatic stress disorder, sleep, polysomnography, quantitative electroencephalography, spectral analysis

## Abstract

**BACKGROUND:** Sleep problems are a core feature of post-traumatic stress disorder (PTSD). However, a robust objective measure for the sleep disturbance in patients has yet to be found.

**METHODS:** The current study assessed EEG power across a wide frequency range and multiple scalp locations, in matched trauma-exposed individuals with and without PTSD, during rapid eye movement (REM) and non-REM (NREM) sleep. In addition, a full polysomnographical evaluation was performed, including sleep staging and assessment of respiratory function, limb movements and heart rate. The occurrence of sleep disorders was also assessed.

**RESULTS:** In PTSD patients, NREM sleep shows a substantial loss of slow oscillation power and increased higher frequency activity compared to controls. The change is most pronounced in right-frontal brain areas and correlates with insomnia. PTSD REM sleep shows a large power shift in the opposite direction, with increased slow oscillation power in occipital areas, which is strongly related to nightmare activity and to lesser extent with insomnia. These pronounced spectral changes occur in the context of severe subjective sleep problems, increased occurrence of various sleep disorders and modest changes in sleep macrostructure.

**CONCLUSIONS:** This is the first study to show pronounced changes in EEG spectral topologies during both NREM and REM sleep in PTSD. Importantly, the observed power changes reflect the hallmarks of PTSD sleep problems: insomnia and nightmares and may thus be specific for PTSD. A spectral index derived from these data distinguishes patients from controls with high effect size, bearing promise as a candidate biomarker.

## Introduction

Posttraumatic stress disorder (PTSD) is a highly debilitating disorder with a lifetime prevalence of 7-8% (1,2). Sleep problems are the most prevalent symptoms of PTSD with roughly 70% of patients suffering from co-occurring sleep disorders (3,4). The sleep problems typically include nightmares, distressed awakenings, nocturnal panic attacks, sleep terrors and insomnia (5).

Sleep abnormalities following trauma are a strong predictor for future development of PTSD (6–9). However, sleep abnormalities may also pre-date trauma (10) and then also predict subsequent PTSD (9). These and other findings suggest that sleep disturbances play an important role in the development and maintenance of PTSD (11). This role is likely related to sleep’s crucial involvement in memory consolidation (12,13), reduction of memories’ emotional tone (14–17) and emotional regulation in general (14,18,19).

While the subjective sleep impairments in PTSD are well established, the underlying physiological abnormalities are less clear. Several studies have focused on the macrostructure of sleep, measuring variables such as the amount of time slept, periods of wakefulness after sleep-onset and the amount of time spent in the different sleep stages. A meta-analysis of such studies reports an increase in stage 1 sleep (the lightest sleep stage), decreased slow wave sleep (SWS) or deep sleep, and increased density of rapid eye movements (REMs) in PTSD patients (20). The reported average effect sizes are modest, however, with small effects for stage 1 (d=0.24) and SWS (d=-0.28), and a small to medium effect for REM density (d=0.43).

While sleep stages are informative about the temporal organization of sleep, they have limited value for quantifying the spectral content of the sleep EEG. Sleep staging is by definition a categorical analysis in which a large amount of variance with regard to the frequency content of the signal goes undetected. Moreover, sleep staging regards neuronal population dynamics as a whole brain state, resulting in a spatially globalized analysis. However, EEG-recorded activity varies substantially across different regions of the brain, during sleeping and waking alike. This spatial variance is largely disregarded in standard polysomnography.

A more precise way to quantify the frequency content of the EEG concerns the analysis of spectral power. A few studies have used such analyses in controlled studies of PTSD sleep, albeit only on recordings from central derivations (21–25). The most consistent observation in these studies is reduced non-REM (NREM) delta (0.5-4Hz) activity in patients (23,24,25). Activity in this frequency range indexes sleep depth. Thus, the reduction in delta power suggests reduced sleep depth in PTSD, in line with the findings of reduced SWS. However, as with the macroscopic findings, reduced delta activity is not consistently observed in all studies (22,26). Thus, none of the physiological sleep measures assessed thus far provides a reliable correlate of the severe subjective sleep problems in PTSD.

The primary aim of the current study was to obtain a better understanding of the neural underpinnings of PTSD sleep disturbances, through a detailed assessment of spatially distributed brain activity. Secondly, we hoped to find a robust neural correlate of PTSD sleep problems, leading the way toward development of a biomarker. Such a biomarker would importantly facilitate further research on PTSD, including the objective evaluation of (sleep-oriented) therapeutic strategies.

With these goals in mind, we performed a comprehensive investigation of sleep in patients with PTSD and matched, traumatized subjects without PTSD. Central to the investigation was a spectral analysis of the sleep EEG across multiple brain areas. We considered a broad frequency range (0.5-50Hz) and adopted a division into frequency bands in line with state-of-the-art knowledge on the neural underpinnings of oscillatory population dynamics. In particular, we separated the delta band, considered in previous studies (0.5-4Hz), into slow oscillations (~0.5-1.5Hz), which have a cortical origin (27–29), and higher delta frequencies (~1.5-4Hz), which have a thalamic origin (30).

As an *a priori* hypothesis, we considered that slow oscillation (SO) dynamics might be disrupted in PTSD. Indeed, SOs (31) orchestrate sleep-related cortical communication (32), and have been shown to play an important role in (emotional) memory consolidation (33,34) and emotional resilience (14). As such, their disruption might lead to abnormal trauma memory consolidation and impaired recovery from emotional trauma. Furthermore, SO dynamics are most pronounced in EEG recordings over frontal cortical areas. The moderate reductions of 0.5-4Hz power, observed over central locations previously, might actually reflect a SO power deficit, picked up in diluted form.

While the above hypothesis concerns NREM sleep, the subjective sleep complaints associated with PTSD are likely associated to both NREM and REM sleep: insomnia, one of the most common PTSD symptoms, is thought to arise primarily from NREM sleep disruption, as are nocturnal panic attacks and sleep terrors (35,36). On the other hand, nightmares, the other core feature of PTSD sleep, are primarily a REM sleep phenomenon (37) although they may also occur during NREM sleep in patients with PTSD (38). Therefore, we expected to find physiological abnormalities also in REM sleep.

In addition to the EEG spectral analysis, a full sleep physiological evaluation (polysomnography) was performed, including visual sleep staging, assessments of limb movements, respiratory, and cardiac function. Furthermore, we evaluated subjective sleep quality and the presence of sleep disorders by means of validated questionnaires. These measures allowed us to relate any abnormalities in brain activity to sleep symptomatology.

## Method

### Participants

Participants in the experiment were traumatized police officers and military veterans, with (N=16) or without (N=14) PTSD. Participants with PTSD were recruited at Foundation Centrum’45, the Dutch national center for diagnostics and treatment of PTSD, part of Arq Psychotrauma Expert Group. Participants in the PTSD and control group were matched on age, gender, education level and professional background (table 1). Participants were asked to refrain from medication use prior to the experiment. However, in seven PTSD patients medication (SSRI’s, antipsychotics, anti-depressants, sedatives or hypnotics) could, for medical ethical reasons, not be interrupted. All participants gave written informed consent. For more information see the supplementary materials.

**Table 1.** Sociodemographic and clinical characteristics of participants with PTSD and trauma-exposed controls.

| Characteristic | PTSD group | Control group |
| --- | --- | --- |
| *Professional background (n, %)* |  |  |
| Police | 12 (75%) | 10 (71%) |
| Veteran | 4 (25%) | 4 (29%) |
| *Mean age (SD)* | 45.6 (7.9) | 44.4 (8.7) |
| *Gender (n, %)* |  |  |
| Male | 15 (94%) | 13 (93%) |
| Female | 1 (6%) | 1 (7%) |
| *Educational level (n, %)* |  |  |
| Lower vocational education | 2 (14%) | 0 (0%) |
| Middle vocational education | 9 (64%) | 11 (79%) |
| Higher vocational education | 3 (21%) | 3 (21%) |
| *Test scores (mean, SD)* |  |  |
| CAPS score | 82.8 (11.6) | 5.3 (4.7) |

CAPS: clinical-administered PTSD scale.

### Clinical assessments and sleep-related questionnaires

The presence or absence of PTSD according to DSM-IV criteria was diagnosed with the Clinician Administered PTSD Scale (CAPS), following assessment of trauma history with the Life Events Checklist (39). We used the CAPS score on item B2 as an indicator of nightmare severity (combined score of frequency and intensity). Past and present comorbid psychiatric disorders according to DSM-IV criteria were assessed with the MINI-PLUS clinical interview (40).

The presence of sleep disorders that were not physiologically evaluated (see next section) was assessed with the SLEEP-50 (41). Subjective sleep quality on the night of the polysomnography was assessed the morning after, with the Dutch Sleep Quality Scale (42).

### Polysomnography and general procedure

All experimentation, including sleep recording, was performed at the clinical facility of Centrum’45. Subjects slept at the clinic twice in the context of a broad diagnostic assessment, before treatment onset. Data for the current study was recorded on one of the two nights, with the order of the nights counterbalanced over subjects and disease status (PTSD, trauma-control). Subjective sleep quality did not differ significantly between the two nights, either in participants with PTSD or in trauma-controls (p's>0.1). All reported clinical and sleep diagnostics were obtained within six to eight weeks of the sleep recording.

Polysomnographic data was recorded successfully for 13 participants with and 14 without PTSD. Subjects were given the opportunity to sleep undisturbed for 9 hours during a lights-off period starting between 11 and 12 PM, depending on habitual sleep times. Polysomnography, using ambulatory 16-channel Porti amplifiers (TMS-i) and Galaxy sleep analysis software (PHI-international), consisted of EEG recording (F3, F4, C4 & O2, referenced to average mastoids), two EOG electrodes monitoring eye-movements, and two for submental EMG. Further sensors were ECG monitoring heart rate, plethysmography monitoring blood oxygenation, piezo leg sensors to detect leg movements, probes measuring nasal airflow, and a piezo respiratory band for thoracic respiratory effort to monitor breathing and sleep apnea. Sample rate for all signals was 512Hz.

### Data Analysis

Sleep stages were scored visually according to AASM criteria (43). For each recording, we calculated total sleep time, sleep latency, REM latency, time awake after sleep onset, and sleep efficiency. We also determined the amounts of light sleep (N1+N2), SWS (N3) and REM sleep, in minutes and as percentage of total sleep time.

The frequency content of the EEG was analyzed using fast Fourier transform-based spectral analysis (4s time windows with 50% overlap, 0.25Hz bin size; Hamming window), on each electrode (F3, F4, C4 & O2) for NREM sleep and REM sleep separately. For each frequency bin, the power per 30s epoch was computed. Normalized power for each epoch in each sleep stage was calculated dividing power per frequency bin by total power in the 0.5-50Hz range. Finally, normalized power bins were merged across frequencies in each of the following bands: SOs (0.5–1.5Hz), delta (1.5–4Hz), theta (4–8Hz), alpha (8–12Hz), sigma (11–16Hz), beta (12–30Hz) and gamma (30– 50Hz).

Apneas and hypopneas, oxygen desaturations, leg movements and R-peaks in the ECG were automatically scored (Galaxy, PHI-international) and manually checked. From these measures an apnea/hypopnea index, oxygen saturation index, periodic leg movement index, leg movement index and heart rate were calculated (details in Supplementary materials).

Sleep macrostructure and non-EEG physiological variables were statistically analyzed using independent samples t-tests (two-tailed) or Mann-Whitney U tests. Relative spectral power was log-transformed to achieve a Gaussian distribution and analyzed through repeated measures ANOVA with factors Diagnosis (PTSD, trauma-control), Sleep State (NREM, REM) and Electrode (F3, F4, C4, O4). The model contained all main and interaction effects of the factor Diagnosis. Frequency bands were analyzed separately to avoid sphericity violations. Correlation analyses were performed using Pearson or Spearman correlation coefficients. Finally, effect sizes were calculated as Glass' delta (44).

## Results

### Sleep quality and sleep disorders

Means and standard deviations of subjectively assessed sleep quality and sleep disorders are given in table 2. As expected, PTSD patients rated their sleep quality as extremely poor (t=-4.9, p<0.000) and scored very high on insomnia (insomnia: t=9.3, p<0.000) and nightmares (W=142.5, p<0.000) compared to trauma-controls. Furthermore, symptoms of periodic limb movement disorder were increased in patients (t=2.4, p=0.026). Considering diagnostic threshold criteria, 13 PTSD patients out of 16 met criteria for insomnia, 11 for nightmare disorder, 8 for periodic limb movement disorder and 1 for circadian rhythm sleep disorder. In the control sample, the number of participants crossing a diagnostic threshold ranged between 0 and 3 across all scales. Finally, PTSD subjects’ daily functioning complaints associated to sleep problems were much higher than trauma-controls’ (W=105, p<0.000).

**Table 2.** Questionnaire-based sleep variables.

| **Variable** | **Group** | **Mean** | **STD** | **p** |
| --- | --- | --- | --- | --- |
| Sleep quality | PTSD | 5.9 | 3.8 | *** |
|  | CTRL | 11.6 | 2.6 |  |
| Insomnia | PTSD | 23.3 | 4.9 | *** |
|  | CTRL | 10.9 | 1.7 |  |
| Nightmares | PTSD | 3.8 | 2.8 | *** |
|  | CTRL | 0.1 | 0.5 |  |
| PLMD | PTSD | 7.1 | 2.9 | * |
|  | CTRL | 5.1 | 1.3 |  |
| CRSD | PTSD | 5.2 | 2.3 |  |
|  | CTRL | 5.2 | 2.0 |  |
| Sleep walking | PTSD | 3.3 | 0.8 |  |
|  | CTRL | 3.5 | 1.7 |  |
| Daily functioning | PTSD | 22.0 | 4.1 | *** |
|  | CTRL | 9.9 | 2.5 |  |

PLMD: periodic limb movement disorder, CRSD: circadian rhythm sleep disorder
\*: $P < 0.05$
\*\*\*: $P < 0.000$

Non-EEG physiological measures to assess movement and breathing-related sleep disorders (table 3) showed an increased Limb movement index in PTSD subjects compared to trauma-controls (W=158.0, p=0.039). Heart rate and respiratory variables did not differ significantly between groups (p's>0.1), but one PTSD participant was diagnosed with sleep apnea.

**Table 3.** Non-EEG physiological variables.

| **Variable** | **Group** | **Mean** | **STD** | **p** |
| --- | --- | --- | --- | --- |
| Limb movement index | PTSD | 58.1 | 42.9 | * |
|  | CTRL | 33.8 | 13.6 |  |
| Respiratory disturbance index | PTSD | 1.4 | 2.2 |  |
|  | CTRL | 1.6 | 1.4 |  |
| SpO2 desaturation 3% index | PTSD | 3.3 | 2.5 |  |
|  | CTRL | 4.3 | 3.5 |  |
| SpO2 desaturation 4% index | PTSD | 1.4 | 1.1 |  |
|  | CTRL | 1.8 | 2.0 |  |
| Mean SpO2 | PTSD | 94.0 | 1.6 |  |
|  | CTRL | 94.6 | 1.2 |  |

SpO2: blood oxygen saturation level
\*: P<0.05

### Sleep macrostructure

Sleep macrostructural variables (table 4) also differed between groups. Subjects with PTSD displayed significantly more awakenings during sleep, both in terms of the absolute number (t=2.4, p=0.025) and frequency of awakenings (awakenings/TST: t=3.0, p=0.006), increased total time awake after sleep onset (t=2.3, p=0.037), a tendency toward longer sleep latency (W=164.5, p=0.077) and, consequently, reduced sleep efficiency (t=-2.5, p=0.025). Furthermore, the PTSD group showed trend-level changes in sleep stage composition compared to trauma-controls: N1 percentage was somewhat increased (t=2.0, p=0.056), while there was a decrease in N3 percentage (t=-1.9, p=0.067) and time spent in N3 (t=-2.0, p=0.057). Finally, REM latency in the PTSD group was significantly increased (t=2.2, p=0.043). For other variables no significant differences were found (p’s>0.1).

**Table 4.** Sleep macrostructure.

| **Variable** | **Group** | **Mean** | **STD** | **p** |
| --- | --- | --- | --- | --- |
| Total sleep time (min) | PTSD | 393.0 | 82.6 |  |
|  | CTRL | 429.6 | 45.9 |  |
| Sleep latency (min) | PTSD | 21.1 | 24.1 |  |
|  | CTRL | 10.0 | 5.3 |  |
| REM latency (min) | PTSD | 130.8 | 81.0 | * |
|  | CTRL | 79.3 | 36.3 |  |
| Sleep efficiency | PTSD | 82.9 | 11.5 | * |
|  | CTRL | 90.9 | 3.6 |  |
| Wake after sleep onset (min) | PTSD | 56.5 | 34.9 | * |
|  | CTRL | 33.3 | 16.6 |  |
| WASO/TST | PTSD | 0.2 | 0.1 | * |
|  | CTRL | 0.1 | 0.0 |  |
| Awakenings (nr) | PTSD | 31.3 | 8.5 | * |
|  | CTRL | 24.1 | 7.5 |  |
| Awakenings/TST (#/min) | PTSD | 4.9 | 1.7 | ** |
|  | CTRL | 3.4 | 1.0 |  |
| Arousal index (#/min) | PTSD | 23.8 | 6.8 |  |
|  | CTRL | 21.4 | 7.2 |  |
| N1 (min) | PTSD | 67.3 | 26.2 |  |
|  | CTRL | 57.8 | 22.1 |  |
| N1 % | PTSD | 17.3 | 5.5 |  |
|  | CTRL | 13.4 | 4.8 |  |
| N2 (min) | PTSD | 238.0 | 45.6 |  |
|  | CTRL | 248.7 | 21.3 |  |
| N2 % | PTSD | 61.1 | 6.6 |  |
|  | CTRL | 58.3 | 5.7 |  |
| N3 (min) | PTSD | 18.2 | 19.2 |  |
|  | CTRL | 35.5 | 26.1 |  |
| N3 % | PTSD | 4.4 | 4.2 |  |
|  | CTRL | 8.3 | 6.3 |  |
| REM (min) | PTSD | 68.6 | 36.7 |  |
|  | CTRL | 87.6 | 29.3 |  |
| REM % | PTSD | 17.1 | 7.3 |  |
|  | CTRL | 20.0 | 4.6 |  |

WASO: wake after sleep onset, TST: total sleep time, Arousal index: number of arousals / total sleep time.
\*: P<0.05
\*\*: P<0.01

### Spectral Analysis

The sleep EEG spectral analysis revealed a striking pattern of abnormalities in PTSD patients. An overview of these abnormalities, across the space-frequency domain, is presented in figure 1, which shows, for the PTSD group, the power deviations from control group values (individual group means and SDs are given in supplementary table x). As can be seen, there is a selective loss of SO power in PTSD NREM sleep and a power increase across higher frequency bands (Fig. 1a). The pattern is apparent across all derivations, but is most pronounced on frontal electrodes, especially in the right hemisphere. PTSD REM sleep (Fig. 1b) shows a more or less opposite pattern, with increased SO power and power loss in higher frequency bands. This pattern is most pronounced occipitally.

**Figure 1.**
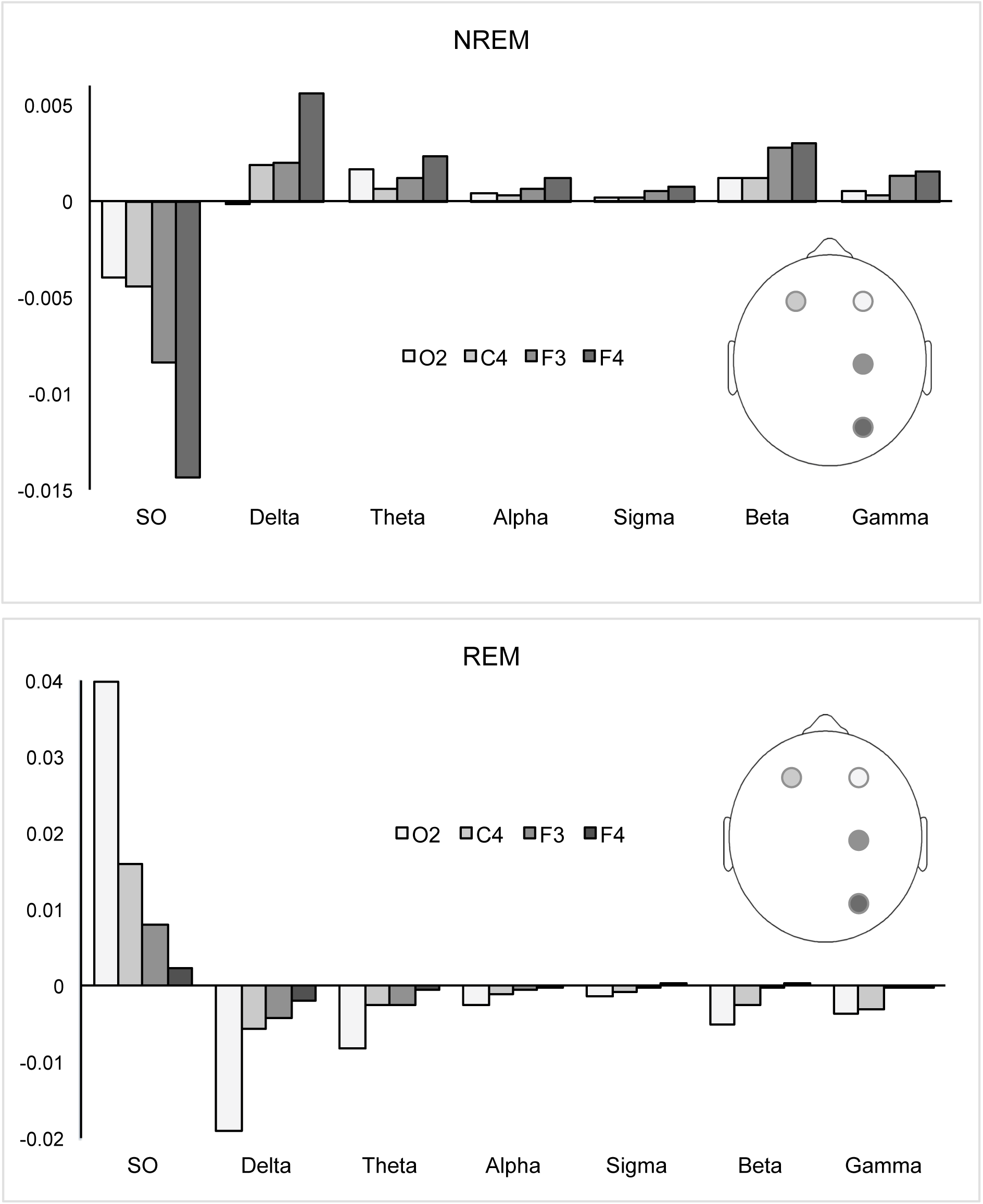
Mean differences in normalized oscillatory frequency power between PTSD patients and trauma-controls in NREM (upper panel) and REM sleep (lower panel), for each derivation and frequency band. Difference values were calculated as [Mean power value PTSD group - Mean power value control group]. SO: slow oscillations.

To provide statistical support for the differential spectral abnormalities in PTSD NREM and REM sleep, repeated measures ANOVA, with factors Diagnosis (PTSD, trauma-control), Sleep State (NREM, REM) and Electrode (F3, F4, C4, O4) was performed for each frequency band. The Diagnosis by Sleep State interaction was highly significant for all frequency bands, with p-values ranging between p=0.001(in the SO band; F=15.2, df1,25, p=0.001) and p=0.008 (gamma band (F=8.2, df1,25, p=0.008). Further analyses were thus conducted for the two sleep states separately, through repeated measures ANOVA with factors Diagnosis (PTSD, trauma-control) and Electrode (F3, F4, C4, O4) (table 5).

**Table 5.** Results of repeated measures ANOVA’s per sleep state and frequency band.

| Frequency band | Effect | NREM |  |  | REM |  |  |
| --- | --- | --- | --- | --- | --- | --- | --- |
|  |  | F | df | p | F | df | p |
| **Slow Waves 0.5-1.5 Hz** | Group | 9.3 | 1,25 | 0.005* | 7.4 | 1,25 | 0.01* |
|  | Group * Electrode | 1.4 | 1,75 | 0.25 | 3.3 | 1,75 | 0.025* |
| **Delta 1.5-4 Hz** | Group | 6.2 | 1,25 | 0.019* | 6.8 | 1,25 | 0.015* |
|  | Group * Electrode | 2.2 | 1,75 | 0.01 | 0.5 | 1,75 | 0.66 |
| **Theta 4-8 Hz** | Group | 4.7 | 1,25 | 0.040* | 5.1 | 1,25 | 0.033* |
|  | Group * Electrode | 1.8 | 1,75 | 0.16 | 0.6 | 1,75 | 0.63 |
| **Alpha 8-12 Hz** | Group | 6.7 | 1,25 | 0.016* | 3.8 | 1,25 | 0.061 |
|  | Group * Electrode | 2.5 | 1,75 | 0.088 | 0.5 | 1,75 | 0.65 |
| **Sigma 12-16 Hz** | Group | 7.0 | 1,25 | 0.014* | 3.7 | 1,25 | 0.065 |
|  | Group * Electrode | 2.4 | 1,75 | 0.08 | 0.6 | 1,75 | 0.62 |
| **Beta 16-30 Hz** | Group | 7.5 | 1,25 | 0.011* | 2.9 | 1,25 | 0.10 |
|  | Group * Electrode | 2.0 | 1,75 | 0.12 | 0.6 | 1,75 | 0.65 |
| **Gamma 30-50 Hz** | Group | 5.3 | 1,25 | 0.030* | 3.7 | 1,25 | 0.066 |
|  | Group * Electrode | 1.5 | 1,75 | 0.23 | 0.5 | 1,75 | 0.70 |

\*: P<0.05

For NREM sleep, a significant effect of Diagnosis was found for all frequency bands (p’s<0.05), supporting that in PTSD power is consistently decreased in the SO range and significantly increased in all other bands, with respect to control. The anterior-posterior gradient in the power change was tested through the Diagnosis*Electrode interaction, assessing the contrast between anterior F4 and posterior O2. The contrast was statistically significant for the delta, alpha, sigma and beta bands (p<0.05) and reached trend-level significance (p<0.1) for all other bands (SO, theta, beta and gamma), confirming larger changes in anterior than posterior region.

For REM sleep, the main effect of Diagnosis was significant for the SO, delta and theta bands and reached trend-level significance in all remaining bands (alpha, sigma, beta, gamma). These results reflect the power increase in the SO band and decrease for higher frequencies in PTSD compared to control. The posterior-anterior gradient in this effect was again assessed through the Diagnosis*Electrode O2 to F4 contrast. The contrast was only significant for the SO band, suggesting that only the SO power increase is significantly localized to posterior brain areas.

Finally, we turned our attention to the spatial locations showing the largest power shifts in each sleep state; that is, right-frontal F4 for NREM sleep and occipital O2 for REM sleep, to assess whether group differences would be statistically robust at the single electrode level. For NREM-F4, the difference between PTSD and trauma-control subjects was highly significant at each frequency band (SO: t=-3.0, p=0.006; delta: t=3.2, p=0.005; theta: t=2.8, p=0.01; alpha: t=3.4, p=0.003; sigma: t=3.3, p=0.004; beta: t=3.3, p=0.003; gamma: t=2.8, p=0.01). Group differences for REM-O2 were slightly less robust, but still reached statistical significance for most frequency bands (SO: t=2.8, p=0.014; delta: t=-2.7, p=0.014; theta: t=-2.6, p=0.018; alpha: t=-2.2, p=0.04, sigma: t=-2.1, p=0.04, beta: t=-1.9, p=0.072; gamma: t=-2.0, p=0.061).

### Correlation of power changes in PTSD with experienced sleep problems

To investigate the relation of abnormalities in oscillatory sleep dynamics with experienced sleep problems in PTSD, we considered the largest power changes in the investigated space-frequency domain. That is, SO power, the most strongly affected band across the combined sleep states, on right-frontal F4 for NREM sleep and on occipital O2 for REM sleep. Each variable was correlated with the two hallmark sleep problems in PTSD: insomnia and nightmares.

Reduced right-frontal SO power in NREM sleep was related to increased insomnia (r=-0.46, p=0.017), but was not related at all to nightmare severity (p>0.1). Conversely, occipital SO power in REM sleep showed an extremely large, highly significant positive correlation with nightmare severity (r=0.84, p=0.003), as well as a significant correlation with insomnia (r=0.46, p=0.016).

### A candidate biomarker for PTSD-sleep problems

We explored the extent to which a single variable, reflecting both the NREM and REM sleep spectral abnormalities, might be used as a biomarker.

To this purpose, we calculated a ‘PTSD spectral sleep index’ (PSSI) as the ratio between right-frontal NREM SO power and occipital REM sleep SO power. The clinical relevance of this index was assessed through the effect size of disease status (PTSD vs trauma-control). We observed an effect size of 3.4, which is considered extremely large (45,46).

Importantly, a biomarker should correlate with diagnostic measures obtained with standardized diagnostic instruments. Accordingly, the PSSI shows a large, highly significant correlation with subjects’ CAPS scores (r=-0.60, p=0.001).

## Discussion

The present study was the first to investigate EEG power in PTSD, over a large space-frequency domain, in both REM and NREM sleep. Our findings reveal substantial power differences between patients with PTSD and traumatized controls. Specifically, patients show a strong shift away from the lowest frequency band (SO band) toward the higher frequencies during NREM sleep, in particular over the right-frontal cortex. On the other hand, during REM sleep SO power is increased at the expense of higher frequency power, in the occipital part of the brain. The latter abnormality is strongly related to nightmare activity, while both REM and NREM abnormalities show a robust relation to insomnia. Abnormalities in PTSD sleep macrostructure were also observed, but were much less pronounced than the power abnormalities. These changes in sleep macro- and microstructure co-occur with severe sleep pathology, apparent from subjective as well as physiological assessments.

The findings in NREM sleep support the hypothesis of deregulated SO dynamics in PTSD, showing a preferential reduction of SO power in patients over frontal areas, where SOs are most frequently generated. Notably, slow oscillations have been implicated in orchestrating sleep-related information processing (32), including memory reactivation and consolidation (47–52). Our findings, therefore, suggest a deregulation of these processes in PTSD. This notion is strengthened by the increased activity in the spindle and beta/gamma bands, which are more directly associated to memory reprocessing (53), thus suggesting exaggerated reprocessing and consolidation in PTSD. Interestingly, key symptoms of PTSD are overgeneralization and aversive intrusion of trauma memories, the latter through flashbacks and nightmares (54,55). These symptoms are reminiscent of runaway consolidation, in which a memory representation is repeatedly and inappropriately reactivated, leading to formation of a giant attractor that dominates network function (56–59). The risk of runaway consolidation is especially high for memories that are strongly encoded to begin with, which certainly applies to trauma memories (detailed account in supplementary material).

The NREM power changes in PTSD patients are larger over right than left prefrontal cortex. This might be related to right hemispheric dominance when it comes to processing emotional or self-relevant information (60,61). Our findings might thus reflect a predominantly right frontal, excessive engagement in the reprocessing and consolidation of traumatic memories in these patients.

Importantly, the frontal loss of SO power also entails a reduction in sleep depth. Sleep depth is traditionally indexed by low frequency power over central electrodes, but recent studies have shown that SO power as well as other oscillatory brain dynamics are also regulated locally, in relation with brain regions’ day-time activity. Deep sleep is essential for sleep's homeostatic recovery function and the resolution of sleep pressure (62). It is, moreover, crucial for a myriad of anabolic and restorative processes, including build-up of energy molecules and the immune system (63–65). Thus, chronic deficiencies in deep sleep have important health consequences (66–68).

Turning to REM sleep, we surprisingly found a power shift toward the SO band, which was most prominent over occipital cortex. The abnormality is highly correlated to nightmare severity. Previous studies have reported increased REM sleep delta (0.5-4Hz) in nightmare disorder (69), and in association with sleep onset hypnagogic imagery (70). Furthermore, delta power is higher in phasic compared to tonic REM sleep (71); the former being the part of REM sleep marked by REMs and hosting most active dreaming (72). The combined findings suggest that increased REM sleep SO power in PTSD patients is related to the frequent nightmares and/or co-concurrently increased REMs. The occipital focus of the power increase, suggests a possible origin in visual cortex involved in dream imagery.

Of note, REM sleep has also been related to memory reprocessing, in particular regarding emotional memories (14,73–75). A comprehensive account of how observed physiological changes during REM and NREM sleep in PTSD might lead to erratic memory consolidation with ‘runaway’ characteristics, is given in a recent review (76).

Our findings regarding sleep macrostructure in PTSD are generally in line with previous findings (20); we observed a notable decrease in sleep efficiency, with increased awakenings and wake-time after sleep onset and a tendency toward longer sleep latency. Patients also showed trend-level changes in sleep stage composition, involving decreased N3 and increased N1, as well as increased REM latency, which was not reported previously. Of interest is a comparison of these results with those of spectral analysis. In fact, the pronounced spectral abnormalities in patients’ sleep EEG are only marginally (for NREM sleep) or not at all (for REM sleep) captured by the analysis of sleep stage composition. Given the nature and limitations of the method, this should not be surprising. Nevertheless, many clinical studies assessing brain activity during sleep still revert to sleep staging as the method of choice. We would like to advocate that spectral analysis across multiple brain locations presents a more sensitive and a more generic method to assess oscillatory brain activity during sleep.

The pronounced spectral abnormalities in PTSD sleep, described above, occur in the context of pronounced sleep pathology. As expected, the large majority of PTSD participants suffered from severe insomnia (81%) and nightmare pathology (69%). More interestingly, we observed excessive limb movements during sleep in a large percentage of patients (50% reached the diagnostic threshold for periodic limb movement disorder). These findings might, in part, be related to the reduced NREM sleep depth in patients, as movement normally diminishes with sleep depth. However, the polysomnographical recordings showed that limb movements also often occurred during REM sleep. This is highly abnormal, as REM-associated hypotonia normally prevents such movements. Indeed, this observation points in the direction of REM sleep behavior disorder, symptoms of which, such as acting out dreams, have clinically been noted in patients with PTSD (77). Interestingly, enhanced leg movements in both REM and NREM sleep have also been observed in nightmare sufferers with and without post-traumatic stress disorder (78). Therefore, the REM sleep movements might be related to the intense negative dreaming and related high arousal, which is experienced by PTSD patients and nightmare sufferers alike.

Besides contributing to our understanding of sleep disturbance in PTSD, our findings may have practical implications. Indeed, the spectral fingerprint of PTSD sleep presents a robust pattern of abnormalities that has not been observed before. These abnormalities, moreover, appear to reflect the main proponents of PTSD sleep pathology, namely insomnia and nightmares, the combination of which has some specificity for PTSD. This fosters the exciting notion that a spectral biomarker of PTSD sleep problems might be obtained. A reliable biomarker would be highly useful in PTSD diagnostic practice and would importantly facilitate further research. Our first steps towards exploring this idea are encouraging, showing that a combined NREM-REM spectral index distinguishes PSTD patients from trauma-controls with strikingly high effect size. This candidate biomarker thus merits further investigation, for instance regarding its specificity versus other sleep and affective disorders.

As a final consideration, and limiting generalization of our results, the patients in this study suffered from severe and chronic PTSD. As such, physiological abnormalities might be particularly pronounced. Also, as the sample consisted of treatment-seeking police officers and veterans, some caution is warranted in extrapolating to other PTSD populations.

In conclusion, our findings reveal substantial, spatially differentiated, abnormalities in the microarchitecture of PTSD sleep, including altered SO dynamics in both sleep states. These changes are likely to affect sleep's homeostatic recovery function and memory reprocessing during sleep. The right frontal hotspot of abnormalities during NREM sleep may be related to reprocessing of negative memories, while the occipital REM sleep abnormalities are related to dysphoric oneiric activity. These findings provide new insights into the neural basis of PTSD-related sleep disorder and its role in PTSD etiology. Furthermore, given their robustness and potential specificity, the sleep microstructural deficits bear promise of delivering a biomarker. Neuroscientifically-informed treatment interventions aimed at targeting specific PTSD symptoms will be essential to future research agendas (79). The objective disease marker for typical sleep problems in PTSD, found in the present study, could importantly enhance such interventions. In particular, those aimed at the debilitating sleep problems associated to PTSD.

## Acknowledgements

We thank Amsterdam Brain and Cognition (ABC), University of Amsterdam, for financially supporting this work. We also thank the staff of Centrum ‘45 for their involvement in the patient recruitment and varied practical support that has been crucial to the realization of this study.

## Financial disclosure section

Dr. M. de Boer reported no biomedical financial interests or potential conflicts of interest.

Dr. M.J. Nijdam reported no biomedical financial interests or potential conflicts of interest.

Dr. R.A. Jongedijk reported no biomedical financial interests or potential conflicts of interest. Prof.

Dr. M. Olff reported no biomedical financial interests or potential conflicts of interest.

Dr. W.F. Hofman reported no biomedical financial interests or potential conflicts of interest.

Dr. L.M. Talamini reported her filed patent on a ‘Device for detecting post-traumatic stress disorder (PTSD) in a subject’ (P100273NL00) and no other biomedical financial interests or potential conflicts of interest.

